# Mechanical Stimuli Affect *E. Coli* Heat Stable Enterotoxin (ST)-Cyclic GMP Signaling in a Human Enteroid Intestine-Chip Diarrhea Model

**DOI:** 10.1101/772608

**Authors:** Laxmi Sunuwar, Jianyi Yin, Magdalena Kasendra, Katia Karalis, James Kaper, James Fleckenstein, Mark Donowitz

## Abstract

Modeling host-pathogen interactions with human intestinal epithelia using enteroid monolayers on permeable supports (such as Transwells) represents an alternative to animal studies or use of colon cancer-derived cell lines. However, the static monolayer model does not expose epithelial cells to mechanical forces normally present in the intestine, including luminal flow and serosal blood flow (shear force) or peristaltic forces. To determine the contribution of mechanical forces in the functional response of human small intestine to a pathogen virulence factor, human jejunal enteroids were cultured as monolayers in microengineered fluidic-based Organ-Chips (Intestine-Chips), exposed to enterotoxigenic *E. coli* heat-stable enterotoxin A (ST), and evaluated under conditions of static fluid, apical and basolateral flow, and flow plus repetitive stretch. Application of flow increased epithelial cell height, transcription of the cyclic nucleotide transporting protein MRP4, and apical and basolateral secretion of cGMP under baseline, unstimulated conditions. Addition of ST under flow conditions increased apical and basolateral secretion of cGMP relative to static conditions, but did not enhance intracellular cGMP accumulation. Cyclic stretch did not have any significant effect beyond that contributed by flow. This study demonstrates that fluid flow application initiates changes in intestinal epithelial cell characteristics relative to static culture conditions under both baseline conditions and with exposure to ST enterotoxin, and suggests that further investigations of application of these mechanical forces will provide insights into physiology and pathophysiology that more closely resembles intact intestine than study under static conditions.

## Introduction

Human adult intestinal stem cell-derived enteroids/organoids retain intestinal segment-specific transcriptional as well as phenotypic characteristics, that enable study of untransformed, non-cancer epithelia (1–4). Enteroids are generally propagated as 3-dimensional (3D) basement membrane matrix-embedded spheroids that are polarized such that the inward facing apical cell surface is not directly accessible. This places limitations on host-pathogen interaction studies. Enteroids grown as monolayers on collagen IV-coated permeable supports overcome the challenges of 3D culture, by permitting direct application of commensal organisms, pathogens, toxins, nutrients, electrolytes and drugs to the apical cell surface and enabling subsequent sampling of exported metabolites from both apical and basolateral surfaces (2, 5–8).

Static enteroid monolayers do not, however, recreate mechanical forces acting on intestinal epithelia *in vivo* (4), lacking luminal flow and blood flow (shear stress forces) and the rhythmic contractions that are part of peristalsis. Although multiple efforts have yielded microengineered cell culture systems with flow to recapitulate physical forces of flow and stretch in Organ-on-Chip models (9–12), evidence of discrete advantages associated with inclusion of mechanical forces for modeling host-pathogen interactions within human intestinal epithelia in these systems is not yet clear. In this study, we evaluated the effects of mechanical forces on both baseline characteristics and the responses to heat stable enterotoxin (ST), a virulence factor of enterotoxigenic *E. coli* (ETEC) infection that is associated with a large burden of diarrheal illness in low-middle income countries (13). The model was a micro-engineered polydimethylsiloxane (PDMS)-based Intestine-Chip (10, 14, 15). The chip contains two parallel hollow channels separated by a flexible permeable membrane that enables human intestinal epithelial cells to be cultured in the presence of physiologically-scaled fluid shear stress forces and repetitive lateral mechanical deformation (14, 15).

ETEC infection primarily targets villus enterocytes of the proximal small intestine (16), and thus human jejunal enteroids were used to seed in Intestine-Chips and then allowed to differentiate (17). ST acts by binding to brush border guanylate cyclase C (GCC) and stimulating cGMP synthesis, that ultimately leads to intestinal fluid and electrolyte loss. ST is structurally and functionally related to the endogenously expressed enteric hormones guanylin and uroguanylin, as well as the synthetic peptide linaclotide, which is used to treat chronic constipation and IBS-C (9, 18). Interestingly, linaclotide has been previously reported to induce secretion of cGMP, that has been suggested as playing a role in symptomatic treatment of IBS-C (19). For this reason, apical (AP), basolateral (BL), and intracellular (IC) pools of cGMP were compared following ST exposure in the Intestine-Chip under static, flow, and flow plus rhythmic deformation (stretch) conditions.

## Methods

### Human jejunal enteroid culture and propagation *in vitro*

Enteroids were established from crypts containing stem cells isolated from jejunal biopsies from healthy adult subjects using the protocol of Sato et al (21, 22), with minor modifications as previously described (2). Enteroids were maintained in Matrigel (Corning, Tewksbury, MA) in expansion medium composed of advanced Dulbecco’s modified Eagle medium/F12 (Life Technologies, Carlsbad, CA) containing 100 U/mL penicillin/streptomycin (Quality Biological, Gaithersburg, MD), 10 mmol/L HEPES (Life Technologies), and 1X GlutaMAX (Life Technologies), with 50% Wnt3A conditioned medium (produced by L-Wnt3A cell line, ATCC CRL-2647), 15% R-spondin1–conditioned medium (produced by HEK293T cell line stably expressing mouse R-spondin1; kindly provided by Dr Calvin Kuo, Stanford University, Stanford, CA), 10% Noggin conditioned medium (produced by HEK293T cell line stably expressing mouse Noggin), 1X B27 supplement (Life Technologies), 1 mmol/L N-acetylcysteine (Sigma-Aldrich), 50 ng/mL human epidermal growth factor (Life Technologies), 1 μg/mL (Leu-15) gastrin (AnaSpec, Fremont, CA), 500 nmol/L A83-01 (Tocris, Bristol, United Kingdom), 10 μmol/L SB202190 (Sigma-Aldrich), and 100 μg/mL primocin (InvivoGen, San Diego, CA). Enteroids were cultured in a 5% CO_2_ atmosphere at 37°C and passaged every 7–12 days. Expansion medium was supplemented with 10 μmol/L Y-27632 (Tocris) and 10 μmol/L CHIR99021 (Tocris) during the first 2 days after passaging. Studies were performed with a jeunal enteroid line (passage numbers 30-45).

### Intestine-Chip

Organ-Chips were fabricated from PDMS and assembled as described (14, 23, 24) (Emulate Inc.,Boston, MA). The top (1 mm high × 1 mm wide) and bottom (0.2 mm high x 1 mm wide) channels were separated by a thin (50 μm) flexible, PDMS membrane containing 7 μm diameter pores with 40 μm spacing and surrounded on either side by two vacuum chambers (1 mm high × 300 μm wide). Chips were sterilized with ethyl alcohol (70%) and distilled deionized water. This was followed by activation of the chip according to the manufacturer’s protocol (Emulate, Inc) (15). The chips were then coated with collagen IV solution (Sigma-Aldrich) (200 μg/ml in PBS) and incubated in a humidified 37 °C incubator for 2 hours. Enteroids were isolated from Matrigel by treating with cell recovery solution (Cultrex) at 4 °C for 30 minutes and dissociated with recombinant enzymes (TrypLE, Gibco) to obtain enteroid fragments.

Fragmented enteroids were then suspended in differentiation medium prepared by removing Wnt3A, R-spondin1, and SB202190 in the expansion medium supplemented with Y-27632 and CHIR99021 for the first two days. Approximately 200 fragments/chip were seeded on the upper channel of the Intestine-Chip and incubated overnight without any flow at 37 °C. The following day (D1 post seeding) the enteroids were perfused at 60 μl/h flow rate modeled to mimic the post prandial cardiac output directed to the small intestine (Supplementary Data) until day 5 (D5). For the “flow plus stretch” condition, cyclic membrane deformation (10% strain; 0.15 Hz) was applied. Both of the dynamic components of Intestine-Chip culture, flow and cyclic stretch, plus control of O_2_/CO_2_ were by the use of the Human Emulation System instrumentation (Emulate Inc.) (15).

The enteroid monolayers were confluent on day 5, at which time they were incubated with differentiation media containing IMBX (1mM) (MP, Biomedicals) and vardenafil (1μM) (Sigma-Aldrich) without (control) or with synthetic STp (1nM) (Bachem, Thermofischer) for 6 hours. ST was exposed to the apical channel only. Apical and basolateral samples were collected for six hours and analyzed for cGMP content by ELISA (Enzo Biochem, Inc). For intracellular cGMP determination, cells were lysed with 0.5M HCl (according to the ELISA protocol). Protein concentration was measured using the Pierce BCA Protein Assay (Thermo Fisher).

### Morphological analysis

Enteroids in Intestine-Chips on day 5 post plating were paraformaldehyde-fixed (4%) and permeabilized in 5% (wt/vol) BSA/ 0.1% (vol/vol) Triton-X 100 before being incubated at 4 °C overnight with primary antibodies directed against ZO-1 (1:100) (Life technologies) and sucrase-isomaltase (1:100) (Developmental Studies Hybridoma Bank, University of Iowa). Next day, cells were washed with PBS and incubated with Alexa Fluor-488 and 568 secondary antibodies (1:100), Hoechst 33342 (10ug/ml), Alexa Fluor-647 Phalloidin (1:200) at room temperature for 1 hour. Images were acquired with an upright scanning confocal microscope (Leica SP5 X MP DMI-6000, 25X water objective).

### Quantitative Real-Time Polymerase Chain Reaction

Total RNA was extracted from monolayers in the Intestine-Chip using the PureLink RNA Mini Kit (Life Technologies) according to the manufacturer’s protocol. Complementary DNA was synthesized from 1 to 2μg of RNA using SuperScript VILO Master Mix (Life Technologies). Quantitative real-time polymerase chain reaction (qRT-PCR) was performed using Power SYBR Green Master Mix (Life Technologies) on a QuantStudio 12K Flex real-time PCR system (Applied Biosystems, Foster City, CA). Each sample was studied in triplicate, and 5 ng RNA-equivalent complementary DNA was used for each reaction. The sequences of gene-specific primers were:

MRP4: Forward: GAAGCGCCTGGAATCTACAA, Reverse: AGAGCCCCTGGAGAGAAGAT

MRP5: Forward: CACCATCCACGCCTACAATAAA, Reverse: CACCGCATCGCACACGTA

RN18S: Forward: GCAATTATTCCCCATGAACG, Reverse: GGGACTTAATCAACGCAAGC

The relative fold changes in mRNA levels of MRP4 and MRP5 between static, flow and flow plus stretch were determined using the 2^−ΔΔCT^ method with human *18S* ribosomal RNA as an internal control for normalization (2).

### Statistical Analysis

Data are presented as means ± SEM. Statistical analyses were conducted using the Student’s t test when comparison was between two conditions and One-way ANOVA Tukey’s multiple comparisons test when compared among more than two conditions with P ≤ 0.05 considered statistically significant. Studies were performed using one jejunal enteroid line derived from a normal human subject. Experiments were repeated at least 3 times.

## Results

1. **Shear stress forces and cyclic strain (mechanical stimuli) induce increased extracellular cGMP secretion under baseline conditions in Intestine-Chip.** Human jejunal enteroids were seeded into the top channel of the Intestine-Chip on collagen IV-coated membrane and grown in differentiation medium (DM) for 5 days prior to each experiment. Apical and basolateral effluents (∼360-500 µl) were separately collected during a 6 h period, and the epithelial cells were lysed for cGMP measurement. Under basal conditions, application of single-pass flow or flow plus stretch for 6 hours did not alter intracellular cGMP content relative to the static conditions (**Fig 1a**). However, both apical and basolateral cGMP secretion were significantly increased in response to perfusion compared to the static conditions (**Fig 1b, c**). There was no significant difference in apical and basolateral cGMP secretion between flow alone versus flow plus stretch. Of note, addition of stretch to flow was associated with large variations in apical and basolateral cGMP secretion.
2. **ST increases cGMP content and enhances mechanically stimulated extracellular cGMP secretion in Intestine-Chip.** Compartmentalization was evaluated under static and mechanical force exposed conditions by introducing ST (1 nM) in the static apical medium or the apical perfusate over 6 h. ST significantly increased intracellular cGMP content in flow and flow plus stretch conditions, but the increase was not significant in the static condition (**Fig 2a**). However, the ST-induced increase in cGMP content was not significantly different comparing the static, flow, and flow plus stretch conditions (**Fig 2a&d**). ST significantly increased cGMP secretion apically under flow and flow plus stretch but not under static conditions (**Fig 2b**). The magnitude of the increase was not statistically different between the flow and flow plus stretch conditions (**Figs 2b&d**). Results were somewhat different for ST-induced basolateral secretion. ST significantly increased BL cGMP secretion only under flow plus stretch (**Fig 2c**). The ST-induced increase in basolateral secretion, which was not significant under static conditions, was significantly increased by both flow and flow plus stretch with the magnitude of the increase not significantly different between flow and flow plus stretch (**Fig 2d**). In conclusion, both apical and basolateral cGMP secretion were stimulated by ST [**Fig 2d**; ST effect calculated as ST minus control]. The amount of secretion was similar under both flow and flow plus stretch and exceeded that of static conditions. In contrast, ST exposure increased intracellular cGMP to a surprisingly small but similar extent under static conditions and with application of flow and flow plus stretch. Hence, we hypothesized that both flow and flow plus stretch altered the ability of the enteroid to transport cGMP out of the epithelial cells in such a way that minimized the increase in intracellular cGMP content.
3. **Mechanical stimuli increase jejunal MRP4 mRNA.** Further studies were performed to understand the mechanism of the increased cGMP secretion. We had previously reported that ST-induced cGMP was transported out of the jejunal enteroid by a process inhibited by the MRP inhibitor,MK571 (3). Thus, the effect of flow and flow plus stretch was determined on jejunal MRP4 and MRP5, which are known to transport cyclic nucleotides. Both flow and flow plus stretch increased MRP4 mRNA levels under baseline conditions; however, the increased mRNA failed to reach statistical significance (**Fig 3b**). Importantly, in the presence of ST, flow plus stretch significantly increased MRP4 mRNA. mRNA for MRP5 was not altered under any of the conditions tested (**Fig 3b**). Of note, MRP5 mRNA expression was lower than that of MRP4. Another possible contributing mechanism to changes in MRP4/MRP5 mRNAs was that physical forces increased cGMP via increasing GCC activity. As shown in **Fig 4**, neither flow nor flow plus stretch altered enteroid GCC activity under baseline conditions or after ST exposure.
4. **Epithelia under mechanical stimuli are more columnar than under static conditions.** Intestinal epithelium is exposed to a wide range of mechanical forces in both normal and pathophysiologic states. To determine whether mechanical forces caused structural changes in the enteroids in the Intestine-Chip, heights of the monolayers were determined by confocal microscopy. Both flow and flow plus stretch significantly increased epithelial cell height compared to the height under static conditions (**Figs 5a&b**). Morphologically, XZ dimensions of cells grown under static condition were 10-15 µm tall with cuboidal appearance. Epithelium under flow and flow plus stretch was 20-25 µm in the XZ direction and resembled columnar enterocytes. Of note, the abundant expression of sucrase-isomaltase on the apical surface of these cells (**Fig 5a**) plus the increase in mRNA expression for sucrase-isomaltase and SLC26A3 compared to differentiated enteroids on Transwell filters **(Supplmentary data 2),** suggest that the enteroids cultured on chips are polarized and consist of differentiated enterocytes under all of the applied conditions, despite their different morphologies.

**Figure 1.**
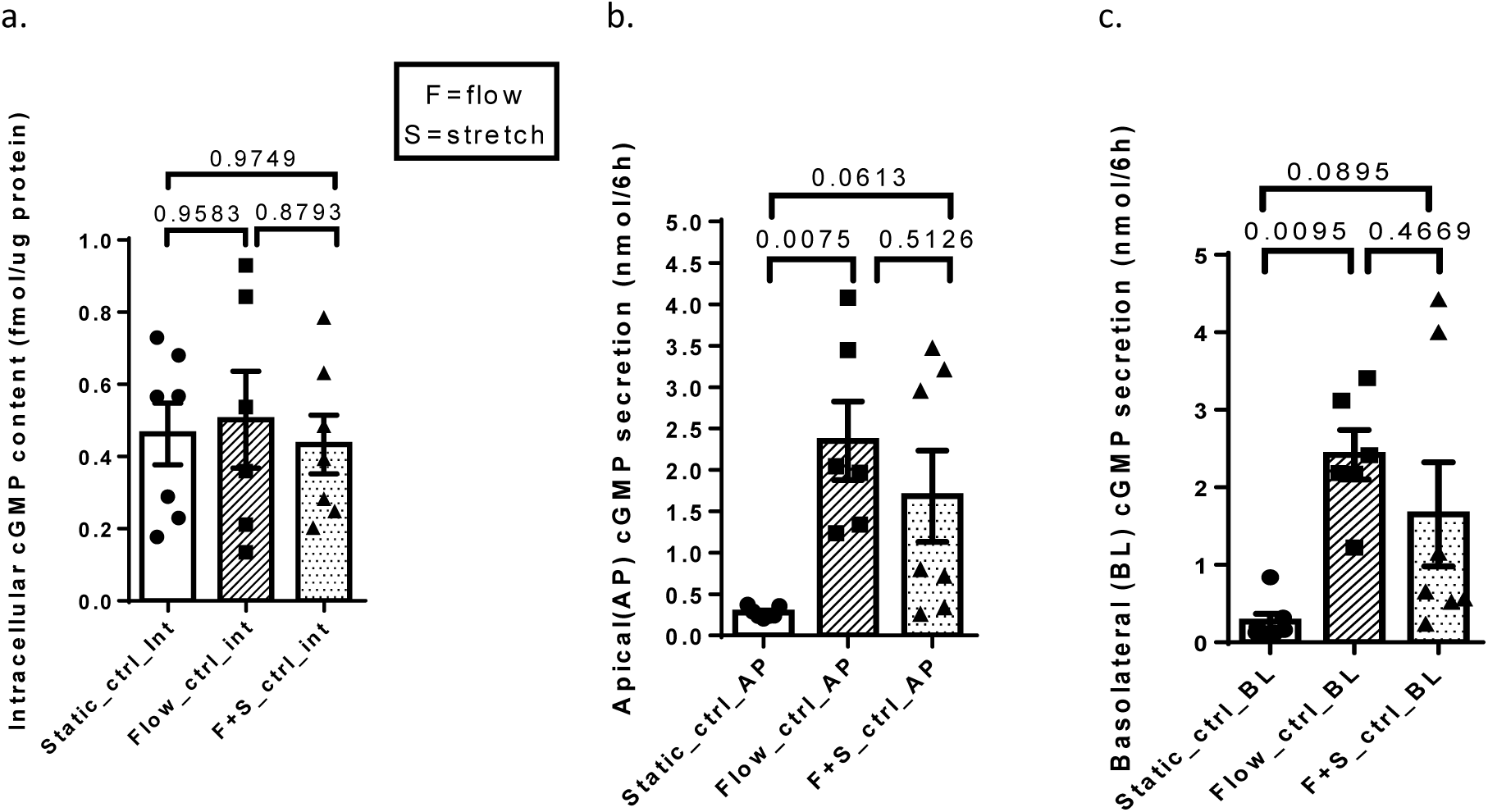
Basal levels of apical (AP) and basolateral (BL) cGMP secretion were increased with shear force provided by flow. (a) Intracellular cGMP content was similar in all conditions. (b&c) AP and BL cGMP secretions were significantly higher with application of sheer force by flow compared to that under static conditions; flow plus stretch caused a similar magnitude of increase but, did not reach statistical significance when compared to static conditions. There was no significant difference in the secretion caused by flow vs flow plus stretch. Static (n=7); flow (n=6); flow plus stretch (n=7). Results are means ±SEM. p values are One-way ANOVA Tukey’s multiple comparisons tests.

**Figure 2.**
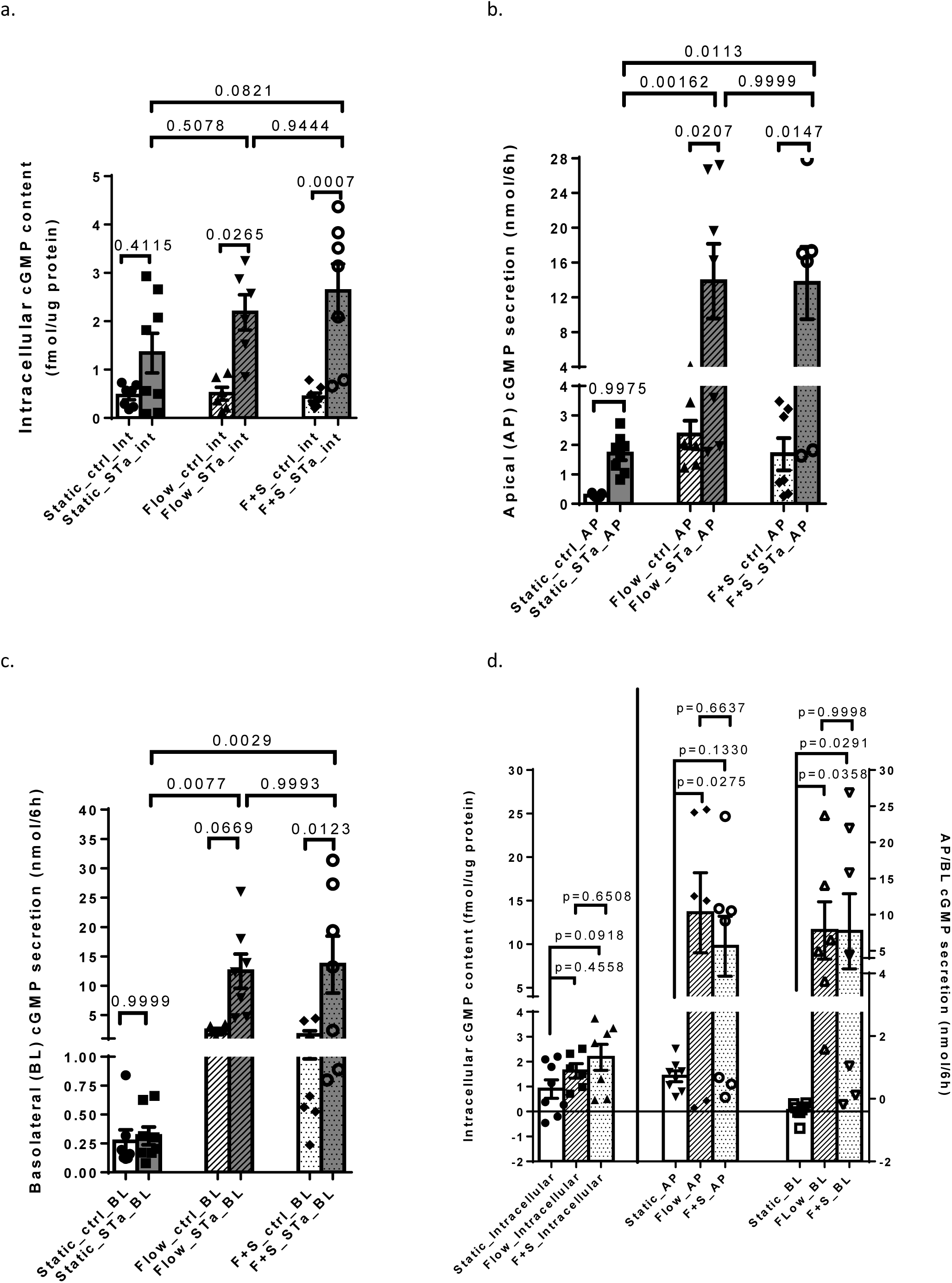
Mechanical stimuli increased ST-induced AP and BL cGMP secretion but did not alter the ST-increase in intracellular cGMP content. (a) ST increased the intracellular cGMP content under all conditions, although the increase was not significant in the static conditions. (b) ST significantly increased AP cGMP secretion with flow and flow plus stretch but, not in the static conditions. There was no difference in the AP cGMP secretion between the flow and flow plus stretch conditions. (c) ST significantly increased BL cGMP secretion only under flow plus stretch condition. However, there was no difference in the BL cGMP secretion between the flow and flow plus stretch conditions. (d) The ST effect on AP cGMP secretion (difference of cGMP secretion in the presence of ST minus that under basal conditions performed on paired samples) under flow and flow plus stretch was significantly greater than the ST effect under static conditions. The ST effect on BL cGMP secretion was significantly higher under flow plus stretch compared to that under static conditions and was increased although not statistically significantly under flow alone. The ST-induced increase of AP and BL cGMP secretion under both flow and flow plus stretch was not different from each other. Static basal (n=7) and ST (n=8); flow basal (n=6) and ST (n=6); flow plus stretch basal (n=7) and ST (n=7). Results are means ±SEM. p values are One-way ANOVA Tukey’s multiple comparisons tests.

**Figure 3.**
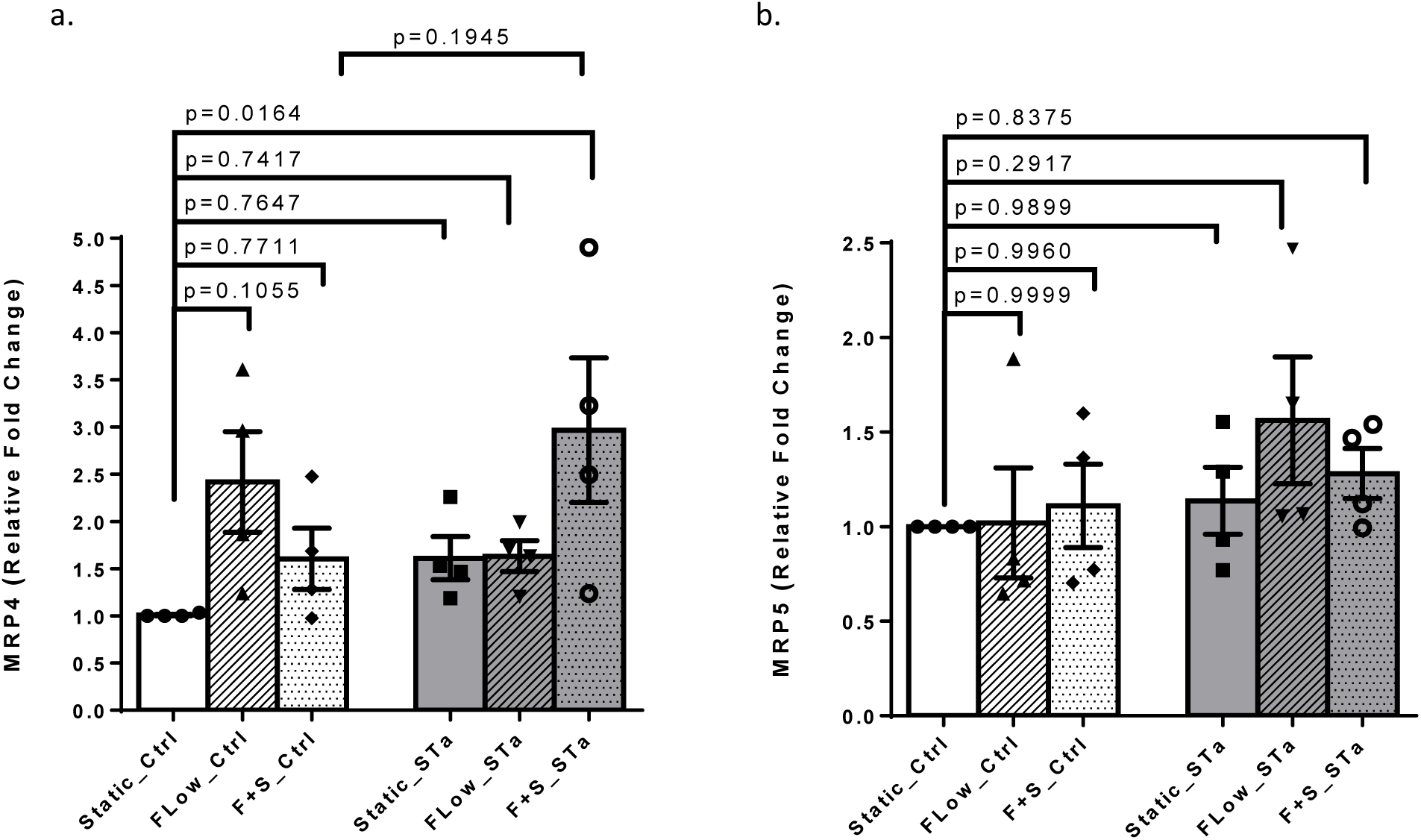
Flow significantly increased MRP4 but not MRP5 mRNA. (a) & (b) MRP4 and MRP5 mRNA were determined by qRT-PCR under basal conditions and after 6 h of ST exposure. (a) Message for MRP4 was increased by flow and flow plus stretch under baseline conditions (statistically not significant), while ST significantly increased MRP4 mRNA under conditions of flow plus stretch. (b) In contrast, MRP5 mRNA was similar under basal conditions and did not change significantly with ST treatment. The level of MPR5 mRNA was lower than that of MRP4. n=4. Results are means ±SEM. p values are One-way ANOVA Tukey’s multiple comparisons tests.

**Fig 4.**
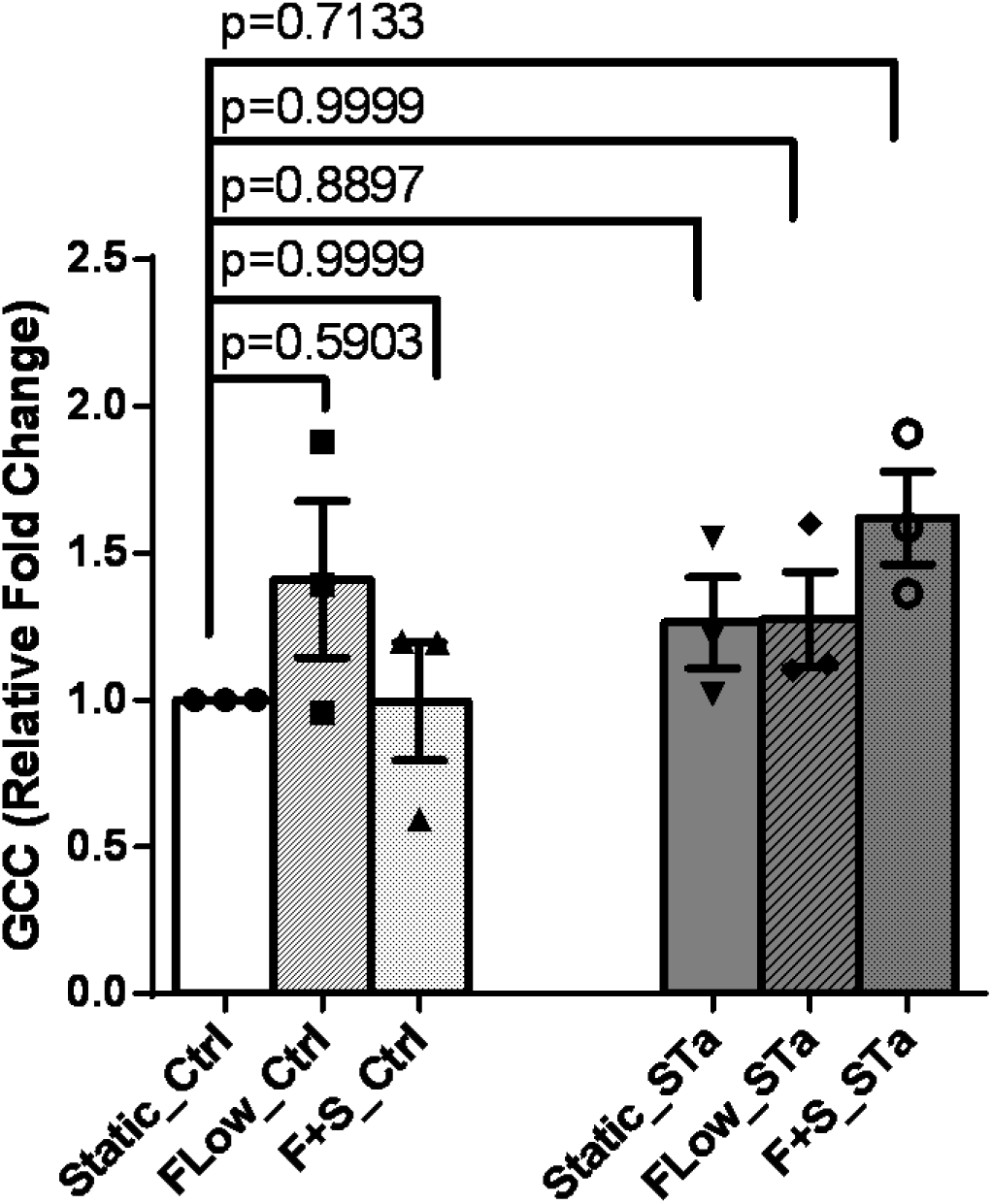
GCC mRNA was not altered by mechanical stimuli. GCC mRNA of enteroids under static, flow and flow plus stretch measured did not show any change either between these three conditions in the basal state or when ST was added. n=3. Results are means ±SEM. p values are One-way ANOVA Tukey’s multiple comparisons tests.

**Figure 5.**
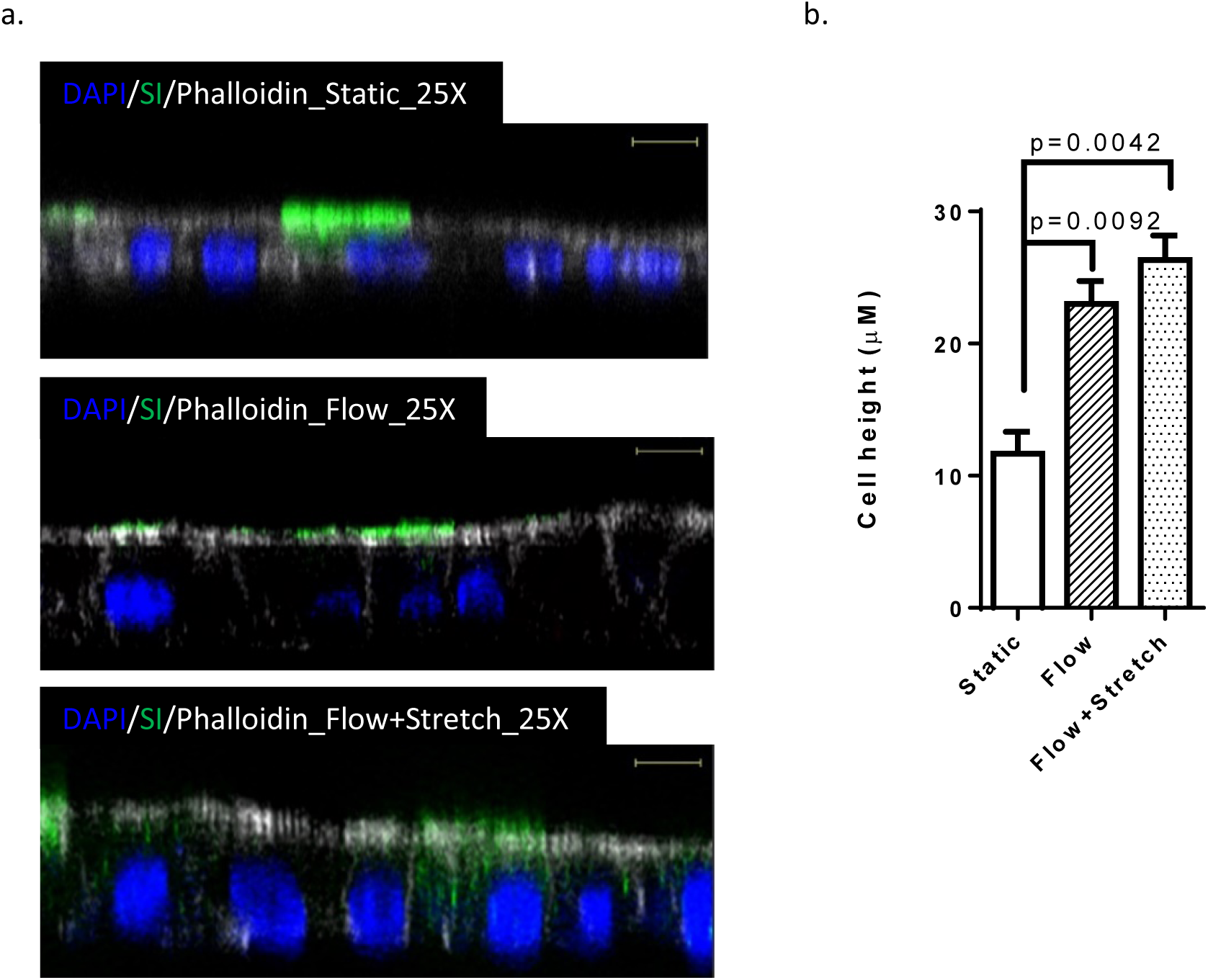
Mechanical stimuli enhance epithelial cell height. (a) Representative XZ views of confocal microscopic images showing jejunal enteroids immunostained for phalloidin (white), sucrase-isomaltase (green) and nucleus (blue) under static (upper), flow (middle) and flow plus stretch (lower) conditions. Bars indicate 10 µm. (b) Quantitation of the cell height under static, flow and flow plus stretch. For all conditions, n=3 regions of interest from two different set of experiments (means ± SEM). p values are Student’s paired t tests.

## Discussion

The mechanical forces exerted on the intestine by luminal flow and blood flow and the cyclic strain of peristalsis have been established as playing an important role in normal physiologic and pathologic states (24–27). These forces affect all of the multiple cell types in the intestine and thus the effects directly on the epithelial cells have been difficult to describe in isolation. The study of human stem cell-derived enteroid monolayers has allowed characterization of normal human epithelial cells. In addition, Ingber, et al. developed a gut-on-a-chip platform they called the “Intestine-Chip” which has allowed application of apical flow that recreates intestinal luminal fluid flow and basolateral flow in the area that blood flow normally occurs (10, 14, 15, 23, 29, 30). Administering repetitive vacuum pressure to mechanically stretch and relax the flexible PDMS membrane and adherent tissues in the central channel, has been applied to mimic peristaltic motions of the living human small intestine. The Intestine-Chip has been shown to alter some structural and functional aspects of human intestinal cells (15, 23, 29); however, most of these studies were performed with the human colon cancer cell line, Caco-2 co-cultured with endothelial cells (15). Here, we extended these studies using the Intestine-Chip and normal human jejunal enteroid-derived chips to examine the pathophysiologic effects of a virulence factor (ST) from a human enteric pathogen, ETEC, in order to begin evaluation of the contributions of these mechanical forces on normal human intestine and in intestinal host-pathogen interactions. In addition, to understand the contribution of the mechanical forces to structure and function of enteroids, we compared the effect of these forces to the same enteroids, which were derived from the same donor and were studied in the absence of the physical forces but in the same platform, referred to as the “static” condition, acknowledging that the Intestine-Chip was designed to include the mechanical forces.

The enteroids were studied in the differentiated state, which we had shown previously to represent the small intestinal villus compartment (2, 17). The concentration of ST studied corresponded to the level of this toxin that was produced when these jejunal enteroid monolayers were exposed to 10^9^ ETEC for 6-8 hours in a Transwell filter (3). The flow rate in the current study was chosen to recapitulate the post prandial cardiac output to the intestinal mucosa [see Supplementary Data 1 (31–33)].

Compared to the static jejunal enteroid monolayers in the Intestine-Chip, flow alone significantly increased the apical and basolateral cGMP secretion under baseline conditions, along with increasing expression of MRP4 mRNA. Morphologically, both flow and flow plus stretch equally increased the epithelial cell height. Flow and flow plus stretch also equivalently increased the ST-induced increases in intracellular cGMP content and apical and basolateral cGMP secretion. However, the ST-induced basolateral cGMP secretion under flow conditions failed to reach statistical significance. Of note, the magnitude of the ST-induced increase in basolateral cGMP secretion was not different when flow and flow plus stretch were compared (**Fig 2d**). MRP4 expression was increased under basal conditions of flow, although ST did not have an additional effect on MRP4 mRNA. This increase in MRP4 potentially contributes to the increased cGMP secretion both under basal and ST exposed conditions. Knockdown studies are being undertaken in a separate study to examine if altering MRP protein expression alters ST-induced cGMP secretion (in preparation by Foulke-Abel, Donowitz). We conclude that the major effect of regulation of cGMP secretion is driven by flow, since addition of stretch did not significantly affect either the intracellular cGMP content or extracellular cGMP secretion beyond that caused by flow alone.

This aim of this study was to characterize the contribution of mechanical forces created by flow and stretch to the effect of a single pathophysiologic enterotoxin (ST) on human enteroids. The data indicates an effect of flow but no additional effect of stretch on ST effects on jejunal cGMP handling. We also point out that multiple factors must be considered in evaluating the relevant normal physiology being modeled in the Intestine-Chip, including: a) The flow in the Intestine-Chip is laminar, while *in vivo* the intestine experiences turbulent flow. This could create some differences in the effects. b) The system of repetitive stretch used was designed to recreate the repetitive distention that occurs with respiration (34), while peristalsis consists of more complex repetitive sweeping cephalad to caudal contractions, that are not entirely regular. Thus, the muscle contractions that occur with peristalsis, provide a more complex force than occurs with the Intestine-Chip. c) Other studies with the Intestine-Chip used one of several endothelial cell types, including intestinal microvascular cells, the effects of which were not considered here (15, 29). This difference may have contributed to the lack of effect of the stretch on the functions studied here.

In the static conditions in the Intestine-Chip, the epithelial cells were shorter and more cuboidal in appearance than the same cells grown as differentiated monolayers on Transwell filters, while the epithelial cell height of differentiated enteroids grown on Transwell filters (8, 35) was similar to those exposed to flow and flow plus stretch in the Intestine-Chip. However, the extent of differentiation of these enteroids in the Intestine-Chip under all conditions compared to the same jejunal enteroids on Transwell inserts, indicate an increased state of differentiation, as judged by level of mRNAs of sucrase-isomaltase and SLC26A3 (downregulated-in-adenoma (DRA)) (Supplementary data 2) and the equally low level of Lgr5 mRNA (data not shown). We present this to indicate the need for further studies to understand how the epithelial height is determined and point out that there are multiple differences in the conditions the enteroids are exposed to in the two platforms, including the materials the cells were grown on, level of oxygenation, hydrostatic pressure from the enclosed Intestine-Chip vs open state of the Transwell, and the fact that the cells on Transwell filters were grown on expansion media before they were switched to differentiation media 5 days before the experiment (2), in contrast to the cells in the current study which were exposed to differentiation media from the time of seeding.

Because of the failure of ST to significantly increase the cGMP content in jejunal enteroid studied in Transwell filters without the presence of phosphodiesterase inhibitors (3), IBMX and the cGMP specific phosphodiesterase-5 inhibitor, vardenafil (36–38) were included in all conditions. In spite of their inclusion, ST failed to alter intracellular cGMP content under static conditions and the significant increase in cGMP content with flow was surprisingly small, which we attribute in part, to the intracellular removal. Flow and flow plus stretch were important for both apical and basolateral cGMP secretion with either small or no increases occurring under static conditions. We can only speculate why the epithelial cells control intracellular cGMP content so carefully, but the well-described role of intracellular cGMP as an inhibitor of proliferation would predict that it is likely to be stringently regulated (39).

The apical and basolateral secretion of cGMP was similar in magnitude in the Intestine-Chip and while it is the control of intracellular cGMP that appears to drive the regulation of cGMP secretion, there is growing evidence of extracellular cGMP having effects on cell physiology and pathophysiology by an auto- or paracrine mechanism. Luminal cGMP inhibits chloride reabsorption in the medullary thick ascending limb of rabbit tubules, and extracellular cGMP inhibits transepithelial sodium transport in porcine renal tubular cells (40). Potential intestinal effects of extracellular cGMP include regulation by apical cGMP of *E. coli* proliferation, release of ST or other ETEC virulence factors and other effects on the intestinal microbiome, and regulation by basolateral cGMP of immune cells in the lamina propria, which are known to affect ETEC (8, 35), and effects on the enteric nervous system (ENS). We have performed preliminary studies evaluating the effect of apical cGMP on ETEC proliferation and ST release in jejunal enteroids with no effect on either parameter found (data not shown). While there is no defined pathophysiologic role as yet for apical cGMP secretion, multiple studies have suggested that cGMP secreted at the basolateral site of enterocytes may affect the enteric nervous system, particularly by inhibiting pain sensation such as the increased visceral sensitivity that is present in IBS-C. Studies of the effects of cGMP on visceral pain sensation have shown that uroguanylin, a naturally occurring intestinal hormone, activates GC-C leading to apical and basolateral secretion of cGMP in Caco-2 cells and in rat colon. In the latter, the cGMP secreted on the basolateral surface inhibits firing of afferent nerves, and cGMP added directly to rat colon inhibited the neural stretch response, as well. In addition, cGMP inhibited murine colon nociception with the effect much greater in mice with chronically induced visceral hypersensitivity (41–43). Most of these studies were undertaken to support the concept that linaclotide, which clinically is an effective laxative, reduces abdominal discomfort associated with IBS-C; with an important preliminary observation that this effect occurs even without the laxative effect occurring (19, 20).

The mechanism leading to secretion of cGMP is speculated to involve the ABC transporter family members, multidrug resistance proteins (MRP). Using MDCK cells, Wijnholds showed that basolaterally localized MRP5 transports nucleotides. He suggested that since MRP4 is closely related to MRP5, it also was likely to also transport nucleotides (44). Moreover, Wielinga and his group showed that MRP4- and MRP5-overexpressing cells, when stimulated with the nitric oxide releasing compound sodium nitroprusside and the adenylate cyclase stimulator forskolin, effluxed more cGMP and cAMP respectively, compared to basal conditions and did so in an ATP-dependent manner (45). He suggested that MRP4 and MRP5 might function as overflow pumps, decreasing steep increases in intracellular cGMP levels. In the current study, under basal conditions without ST exposure, MRP4 mRNA was increased under flow and flow plus stretch (statistically significant) compared to static conditions in agreement with increased extracellular cGMP accumulation. Of note, MRP4 has some increased specificity for cGMP over cAMP (47). Hoque, et. al demonstrated NHERF1 as a major determinant of MRP4 trafficking to apical membranes of mammalian kidney cells. Moreover, he suggested MRP4 may be localized to either apical or basolateral membranes in polarized cells depending on the cell type (47). However, lack of-specificity of the MRP4 and MRP5 antibodies available have prevented localization of these MRPs as well as quantitation by immunoblotting. Further studies are needed to define the role of MRP4 and MRP5 in human jejunal enteroids and to determine whether other cyclic nucleotide transporters might be affected by flow and/or stretch to affect cGMP secretion.

This study demonstrates that the mechanical shear stress forces provided by luminal and/or basolateral flow, compared to studies in the same platform under static conditions, affects multiple aspects of normal human jejunal enteroid structure and function, and also multiple aspects relating to the effects of *E. coli* heat stable enterotoxin on enteroids. The shear force from apical plus basolateral flow induced an increase in altered the cell height, apical and basolateral cGMP secretion under basal conditions (without ST), and altered host-pathogen interactions including the magnitude of the ST-induced apical and basolateral cGMP secretion. Repetitive stretch did not add to these effects. In this study, only the effects of a single virulence factor of one human diarrheal disease were studied. Thus, we suggest that further studies of host-pathogen interactions are necessary to determine the contribution of these physical forces to normal intestinal physiology and pathophysiology, including host-pathogen interactions, as well as potential contributions to studies of drug pharmacokinetics and personalized drug therapy.

## Acknowledgments

This study was partly supported by NIH NIAID PO1AI125181 and NIH NIDDK P30DK089502 and by a grant-in-aid from Emulate, Inc.

## Supplementary 1. Flow rate derivation

Normal cardiac output (CO) = (4-8) L/min

Average CO = 6 L/min

Distribution of CO to superior mesenteric artery (SMA) = 15% of CO

Distribution of CO to intestinal mucosa-submucosa = (50-90) % of CO (average = 70%)

Hence, estimate of blood flow (BF) to intestinal mucosa-submucosa = 6 L/min X 15% X 70%

Microfluidic/micro-human flow = 10^6^

Therefore,

Blood flow to micro intestine mucosa-submucosa = 6 L/min X 15 X 70/10^6^

= 0.0063 L/min

= 0.63 µl/min

In post-prandial state, BF to intestine increases by ∼50%.

Thus, BF to micro intestine mucosa-submucosa = 0.63 – 0.95 µl/min.

**Figure S2.**
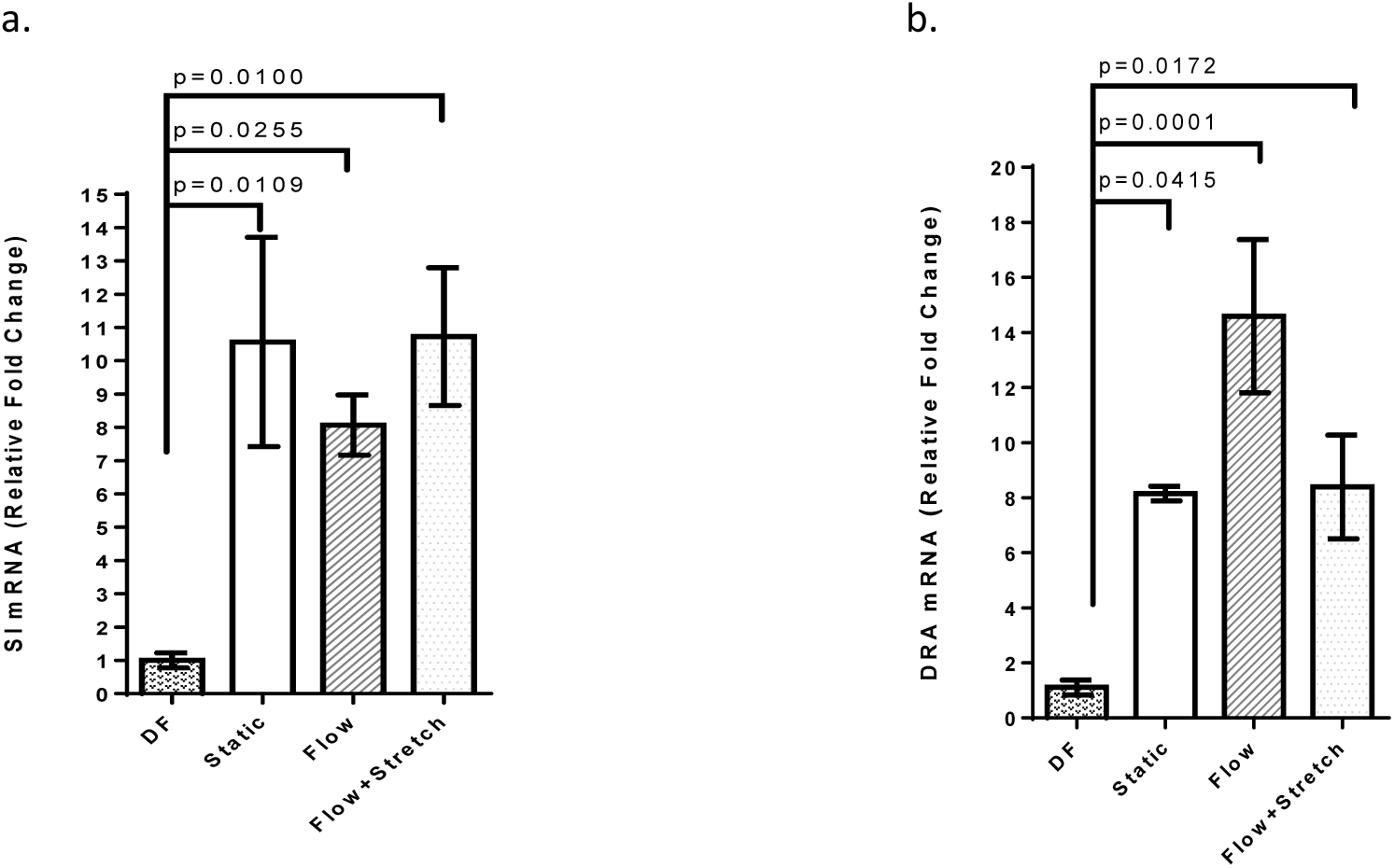
mRNA Differentiation markers SI and DRA were higher in static and flow and flow plus stretch conditions compared to differentiated enteroids in Transwell filters. (a) & (b) Sucrase-isomaltase (SI) and DRA mRNA determined by qRT-PCR in differentiated enteroids in Transwell filters (DF), in Intestine-Chip studied under static, flow and flow plus stretch conditions. mRNAs for both SI and DRA were increased under all conditions in the Intestine-Chip compared to Transwell grown DF enteroids. All conditions in the Intestine-Chip had similar SI mRNAs. For DRA, only the enteroids grown under flow conditions were greater than those grown under static conditions. n=3. Results are means ± SEM. p values are Student’s unpaired t tests.

